# Power spectrum and critical exponents in the 2D stochastic Wilson Cowan model

**DOI:** 10.1101/2022.07.04.498640

**Authors:** I. Apicella, S. Scarpetta, L. de Arcangelis, A. Sarracino, A. de Candia

## Abstract

The power spectrum of brain activity is composed by peaks at characteristic frequencies superimposed to a background that decays as a power law of the frequency, *f^-β^*, with an exponent *β* close to 1 (pink noise). This exponent is predicted to be connected with the exponent *γ* related to the scaling of the average size with the duration of avalanches of activity. “Mean field” models of neural dynamics predict exponents *β* and *γ* equal or near 2 at criticality (brown noise), including the simple branching model and the fully connected stochastic Wilson Cowan model. We here show that a 2D version of the stochastic Wilson Cowan model, where neuron connections decay exponentially with the distance, is characterized by exponents *β* and *γ* markedly different from those of mean field, respectively around 1 and 1.3. The exponents *α* and *τ* of avalanche size and duration distributions, equal to 1.5 and 2 in mean field, decrease respectively to 1.29 ± 0.01 and 1.37 ± 0.01. This seems to suggest the possibility of a different universality class for the model in finite dimension.

## Introduction

In the last two decades, the hypothesis that the brain operates near a critical point has gained a large evidence. The first experiments pointing in this direction were done on organotypic cultures and acute slices of rat cortex [1], where scale-free distributions of activity avalanches were found. Since then, the hypothesis has been confirmed in many systems in vitro and in vivo, from cortical activity of awake monkeys [2] to the resting MEG of human brain [3]. The distribution of avalanche sizes is found to scale as *P*(*S*) ~ *S^-α^*, that of avalanche durations as *P*(*T*) ~ *T^-τ^*, while the mean size of avalanches scales as 〈*S*〉 ~ *T^γ^* as a function of the duration [4–8]. A good indicator of criticality is believed to be given by the scaling relation 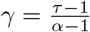, as originally predicted in the theory of crackling noise [9, 10], as well as by the collapse of rescaled shapes of avalanches of different durations [4, 7, 8]. The simple branching model of avalanche propagation predicts exponents *α* = 3/2, *τ* = 2 and *γ* = 2, observed in some experimental realizations [1, 11] and in models of neural dynamics, including the fully-connected stochastic Wilson Cowan model [7]. However, some experimental results have found an exponent *γ* around 1.3, not compatible with the value 2 predicted by the branching process universality class, even when the scaling relation is experimentally satisfied. For instance, results on spike avalanches measured in the urethane-anesthetized rat cortex [12], cultured cortical networks [4], ex-vivo recordings of the turtle visual cortex [5], and somatosensory barrel cortex of the anesthetized rat [13], have found *γ* around 1.3 with the scaling relation 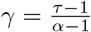 satisfied.

Another important feature of neuronal dynamics is the power law decay *P*(*f*) ~ *f^-β^* of the power spectrum of EEG, MEG, resting state fMRI, and local field potential as a function of frequency [14–18], once the peaks corresponding to characteristic frequencies of oscillations have been subtracted. The values of *β* are found to be between 1 and 1.3 in the EEG and MEG of healthy patients [16, 17], while they are around 2 for epileptic patients [19]. On quite general grounds, the exponent *β* is predicted to be equal to the exponent *γ* of the relation 〈*S*〉 ~ *T^γ^* [9]. It seems therefore that there are at least two universality classes in brain dynamics, one that can be called the “mean field” class, represented by the branching model and the fully connected Wilson Cowan model, that is characterized by *α* ≃ 3/2 and *β* ≃ *γ* ≃ *τ* ≃ 2 (the equality is exact for the branching model), and another characterized by a lower value of the exponents, *β* ≃ *γ* ≲ 1.3, *α* ≃ 1.3, *τ* ≃ 1.4. The experimental measurement of *γ* around 1.3, not compatible with the value of branching process, opened a debate, and it raised the question if a different model, with a different universality class, might be required to explain the critical brain data [12], or if different mechanisms, such as subsampling, should be invoked to reconcile data [20] with branching model.

In the present paper we study the 2D version of the stochastic Wilson Cowan model [7, 21], where neurons are distributed uniformly on a 2D lattice, and connections between them decay exponentially with the distance. We can tune the network topology from 2D to fully-connected by changing the ratio between the range *λ* of the exponential decay of the connections, and the side of the lattice *L*. While in the fully connected case the dynamics of the model is completely described by specifying two variables, the fraction of active excitatory and inhibitory neurons, in the 2D case one needs to define such fractions at each site of the lattice. As a consequence, while the fully connected model is characterized by just two characteristic relaxation times, in the 2D model one finds a whole spectrum of times, related to the different Fourier modes of the neural activity. By making a system size expansion one finds that, if the number of neurons on each site of the lattice is large, the dynamical equations governing the evolutions of the Fourier modes decouple, and each mode obeys to the same equations of the fully connected model, but with a different relaxation time.

Independently of the network topology, the model shows a critical point at a characteristic value of a parameter measuring the difference between the strength of excitatory and inhibitory connections. For values above the critical point, the model displays a self-sustained dynamics even in the absence of external input. At the critical point, one of the characteristic times diverges, related to the mode |***k***| = 0 in spatially extended topologies, while the times of the other modes scale as |***k***|^-2^. This feature, together with the density of the wave numbers which scales as |***k***| in two dimensions, gives rise to a power spectrum that is proportional to *f*^-1^ (pink noise).

In the next section we introduce the model, then we study the power spectrum and relaxation functions of the firing rate in the linear case (large neuron density). We then look at the avalanche size and duration distributions, and show that at the critical point the distributions are scale free, with exponents depending on the topology of the system. Finally, we show that the exponents *β* and *γ* have approximately the same dependence on the inverse frequency and duration of avalanches, respectively, and in the 2D model tend respectively to 1 and 1.3 at low frequency and large durations.

### The model

Let us consider a two-dimensional *L*×*L* lattice, where on each site there are *n_E_* = *N_E_*/*L*^2^ excitatory and *n_I_* = *N_I_*/*L*^2^ inhibitory neurons, with connections depending on the distance rij measured in lattice spacings, as shown in the scheme in Fig. 1. Note that all pairs *i* and *j* of neurons belonging to the same site have distance *r_ij_* = 0, so that inside one lattice site the network is fully connected. Neurons are modeled as in [21], namely they can be in two states, active and quiescent. The rate of transition from active to quiescent state is *α* for all the neurons, while the rate from quiescent to active state is given by an activation function *f* (*s_i_*) of the input *s_i_* of the *i*-th neuron, given by

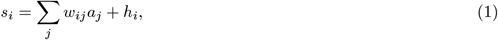

where *a_j_* =0, 1 if the *j*-th neuron is quiescent or active respectively, *w_ij_* are the connections between neurons, and *h_i_* is an external input. We consider the activation function

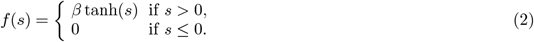

**FIG. 1.**
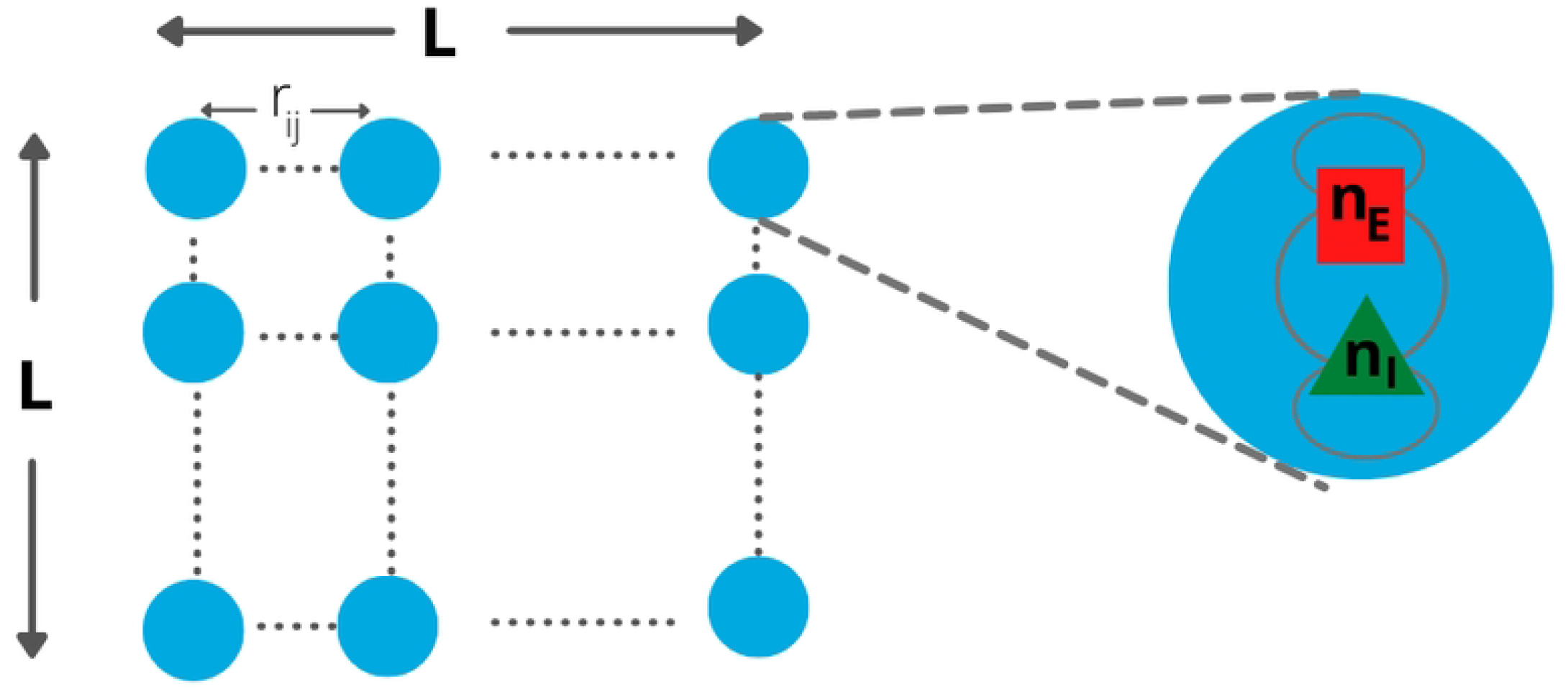
Schematic representation of the 2D model. On each site of the lattice there are *n_E_* excitatory and *n_I_* inhibitory neurons. Connections between neurons depend on the type of neuron (excitatory or inhibitory) and on the distance measured in lattice spacings. Neurons on the same site have *r_i_j* = 0.

In the following we set *α* = 0.1 ms^-1^, and *β* = 1 ms^-1^. We study here a version of the model where the connections between neurons do not depend only on the type of presynaptic neuron, as in the fully connected case, but depend also on the distance between neurons. Namely, the connection between neurons *j* (pre-synaptic) and *i* (post-synaptic) is given by

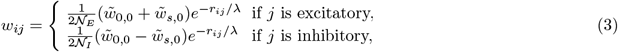

where we have defined 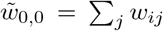 as the sum of all the connections incoming in one neuron, which we take to be the same for all neurons, 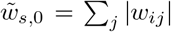, and *r_ij_* is the distance between neurons. The second subscript in 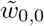 and 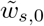 refers to the wave-number ***k*** = 0, see Eq. (13). The normalization factors 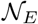 and 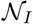 are defined as 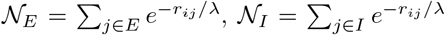, where *E* and *I* are respectively the set of all excitatory and inhibitory neurons. Note that, when *λ* ≫ *L*, we recover the fully connected model, well studied in the past [7, 21, 22]. On the other hand, when *L* ≫ *λ*, the topology of the connections changes to two-dimensional.

We consider the same input *h_i_* ≡ *h* for all neurons. As the connections wij depend only on the distance between neurons, and the input *h* is the same for different neurons, the system is translationally invariant. The fraction Σ of active neurons is a stochastic variable that at stationarity fluctuates around the fixed point value, given by the equation (see Methods)

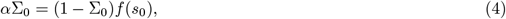

with 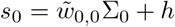. Note that Eq. (4) does not depend on the chosen topology of the connections, fully connected or sparse or depending on the distance, but only on the condition that the sum of incoming connections 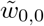 is the same for all neurons, because this makes the system translationally invariant and the fixed point activity Σ_0_ equal for all the lattice sites. In Ref. [7] it was shown that there is a critical point at *h* = 0 and 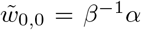. For 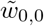 larger than the critical value, an attractive fixed point with Σ_0_ > 0 exists even when the external input *h* → 0. In the case of the fully connected model, the connections wij do not depend on the spatial position of neurons, but only on the functional type (excitatory or inhibitory) of the pre-synaptic and post-synaptic neuron. In Refs. [7, 21, 22] a further simplification was considered, that *w_ij_* depends only on the type of the pre-synaptic neuron, and is given by 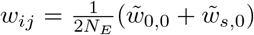 if *j* is excitatory and 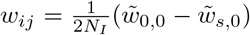 if *j* is inhibitory. In this case, the temporal autocorrelation function of time dependent variables, like the fraction of active neurons or the firing rate, can be written in the limit of large number of neurons as [22]

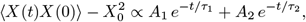

where *X*(*t*) is the variable considered, *X*_0_ is its average value in time, *A*_1_ and *A*_2_ are constant coefficients, and *τ*_1_, *τ*_2_ are the characteristic relaxation times. While *τ*_2_ is always small, and lower than *α*^-1^, the time *τ*_1_ diverges at the critical point [7].

In the fully connected case the state of the system depends only on two variables Σ and Δ, that correspond to the sum and difference between the fractions of activated excitatory and inhibitory neurons, conversely for connections depending on the distance, the values Σ*_**r**_* and Δ*_**r**_* on each site of the lattice are necessary to characterize the activity, defined as

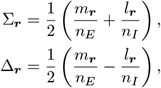

where *m_**r**_* and *l_**r**_* are respectively the number of active excitatory and inhibitory neurons on the lattice site ***r***. Equivalently, we can use the Fourier transforms Σ*_**k**_* and Δ*_**k**_*, where ***k***’s are *L*^2^ different wave vectors. As the system is translationally invariant, the fixed point is characterized by Σ*_**k**_* = Δ*_**k**_* = 0 for ***k*** ≠ 0. Moreover, for the wave vector ***k*** = 0, at the fixed point Δ_0_ = 0 (this is a consequence of the fact that connections do not depend on the post-synaptic neuron), while Σ_0_ obeys the same equation (4) of the fully connected case. Therefore, for 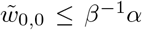, the fixed point value Σ_0_ goes to zero when the external input *h* → 0, while for 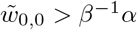 it remains finite even for *h* → 0.

A quantity that will be considered in the following is the local firing rate, defined as

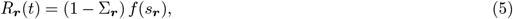

where *s_**r**_* is the input defined by Eq. (1) of neurons on site ***r***. Note that all the neurons on site ***r*** have the same input. The probability that a neuron on site ***r*** fires (makes a transition from inactive to active state) in the interval of time Δ*t* is *R_**r**_*(*t*)Δ*t*.

### Temporal correlations and power spectrum

We now consider a variable *X_**r**_*(*t*) defined on a site ***r***, that can be for instance the fraction of active neurons Σ*_**r**_*(*t*), or the firing rate *R_**r**_*(*t*), given by a superposition of all its Fourier components,

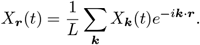

As shown in Methods, for large number of neurons different Fourier components of the activity decouple, and obey the same evolution equation of the total activity in the system with full connectivity. Therefore the autocorrelation function of *X_**r**_*(*t*) will be given by the sum of the autocorrelation functions of its Fourier components,

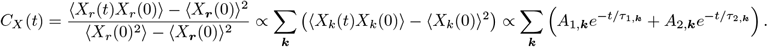

While *A*_2,***k***_ and *τ*_2,***k***_ remain always finite and small, *A*_1,***k***_ and *τ*_1,***k***_ diverge proportionally to |***k***|^-2^ at the critical point. In particular 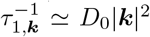, where *D*_0_ is a constant defined in Methods, see Eq. (16). The power spectrum of the activity on a site of the lattice is given by the Wiener-Khinchin theorem as the temporal Fourier transform of the autocorrelation function,

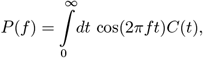

and given the dependence of *A*_1,***k***_ and *τ*_1,***k***_ on the wave number at the critical point, in a system with a two-dimensional structure one finds a dependence *P*(*f*) ∝ 1/*f* for frequencies between 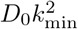 and 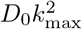, where *k_min_* and *k_max_* are respectively the inverse of the system size and of the lattice constant (see Methods).

In Fig. 2A,B we show the dependence of the autocorrelation function and of the power spectrum on the distance with respect to the critical point, which is given by 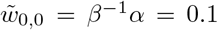 and *h* = 0. Far from the critical point, the distribution of times for different wave vectors is narrow, the autocorrelation decays exponentially with good approximation and the power spectrum can be described by a Lorentzian. Near the critical point, we have a wide distribution of times, that gives rise to a 1/*f* decay in the spectrum. As shown in Fig. 2A, the corresponding relaxation function exhibits the slow decays *a* – *b* ln *t* for a wide interval of times [23].

**FIG. 2.**
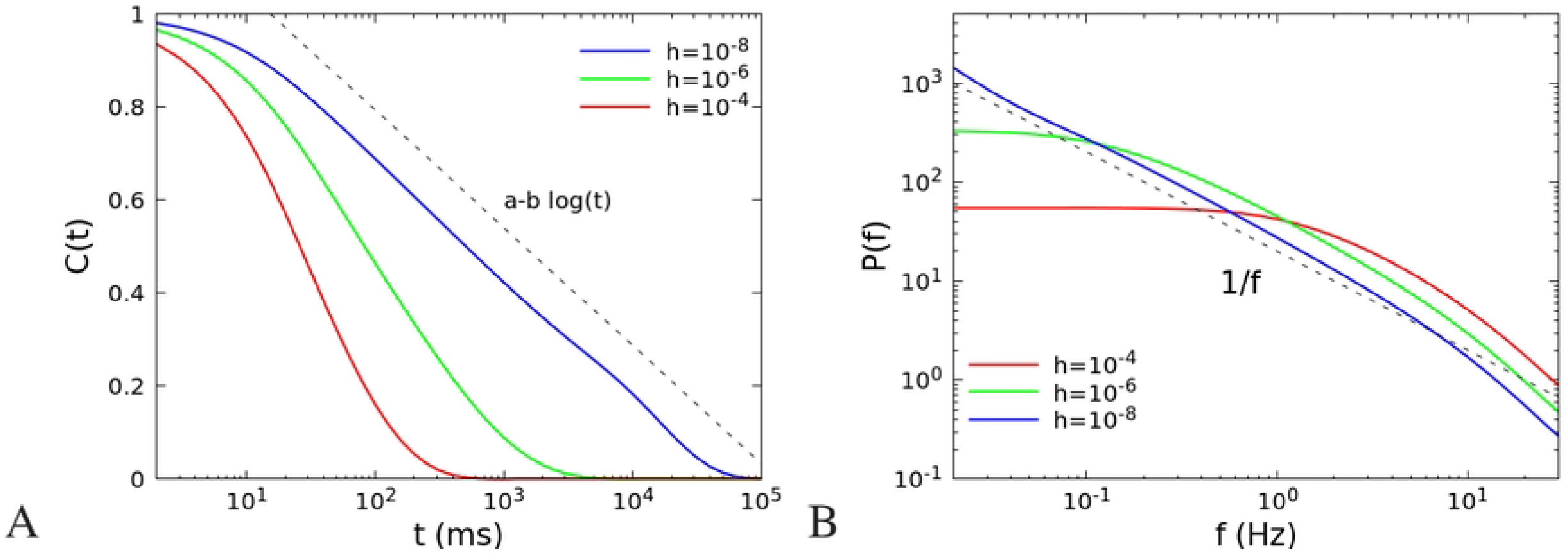
Autocorrelation (A) and power spectrum (B) of the single site firing rate in a two-dimensional *L* × *L* model, in the linear approximation (number of neurons *N* → ∞), as a function of the distance from the critical point. We set *L* = 100, 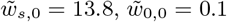, and external input *h* between 10 ^1^ and 10^-8^. The connections between neurons depend on the distance as in Eqs. (3), with *λ* = 1.

As shown in Fig. 3A,B, the 1/*f* dependence of the spectrum depends not only on the distance from the critical point, but also on the range of the connections. If the range *λ* grows, the autocorrelation functions and power spectrum tend to those of the fully connected system, that is characterized by just two correlation times *τ*_1_ and *τ*_2_, where *τ*_2_ is a short time of the order of *α*^-1^ while *τ*_1_ is large near the critical point, so that the autocorrelation is well described at long times by a single exponential.

**FIG. 3.**
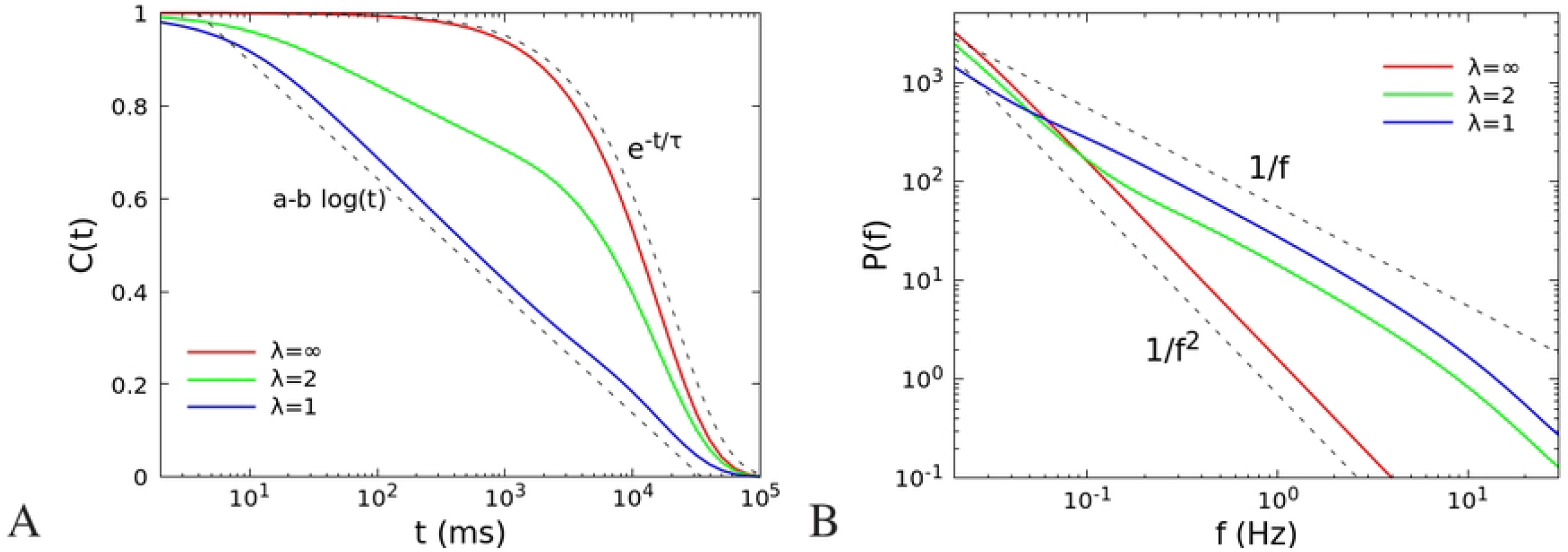
Autocorrelation (A) and power spectrum (B) of the single site firing rate in a two-dimensional *L* × *L* model, in the linear approximation (number of neurons *N* → ∞), as a function of the range *λ* of the interactions. We set *L* = 100, 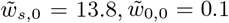, *h* = 10^-8^, and *λ* between 1 and ∞. Note that in the limit *λ* = ∞ the system is no longer 2D but fully connected. The exponent *β* of the power law decay of the power spectrum *P*(*f*) ~ *f^-β^* is *β* = 1 for the 2D model (*λ* = 1, 2) in agreement with the analytical predictions in section Methods, while *β* = 2 in the fully connected case (*λ* = ∞).

### Relation between avalanche and power spectrum exponents

In this section we compare the size and duration distributions of avalanches of activity of a site of the lattice in the 2D case, and in the fully connected model. In the following we set *n_E_* = *n_I_* = 10^8^ neurons for each site, and compare the behaviour of the network with *L* = 40 and *λ* = 1 (2D), with that of a network with *L* =1 (fully connected). We remark here that in a network with *L* = 1 the distance between all neurons is zero, so that in Eq. (3) the connections *w_ij_* depend only on the type of neuron, and the model coincides with the model studied in [7, 21, 22], with full connectivity.

We simulated the system with Langevin dynamics, see Eq. (7). To speed up the simulations, we have set the connections *w_ij_* in Eq. (3) to zero when *r_ij_* > 3*λ*. We made 60 different runs both for *L* =1 and *L* = 40, for respectively 3.5 × 10^7^ ms and 2.5 × 10^5^ ms, discarding the first 10^7^ and 4 × 10^4^ ms, and collecting the avalanches on all sites in the 2D system. We define an avalanche as follow: we divide the time in discrete bins of width *δ* =1 ms, and consider the time evolution of the activity on a single site. We identify an avalanche as a continuous series of time bins in which there is at least one spike (i.e., a transition of one neuron from a quiescent to an active state in the site being considered) [24]. We remark here that avalanches are relative to the activity on a single site, so an avalanche begins and ends when the activity on the considered lattice site is zero, regardless of the activity on the other sites of the lattice. The size of the avalanche is defined as the total number of spikes of neurons belonging to the site considered, while the duration is the number of time bins of the avalanche multiplied by the width δ of the bins. Note that all the lattice sites are equivalent, because the connections are translationally invariant and boundary conditions are periodic, so we expect the same distribution of sizes and durations on all the sites of the network. Therefore, to improve statistics, we compute the average of the distributions over all the sites of the lattice.

The distributions of avalanches size and duration are expected to follow power-law scalings, *P*(*S*) ~ *S^-α^* and *P*(*t*) ~ *T^-τ^*. In Fig. 4 we report the results for the 2D system (*L* = 40, blue dots) and the fully connected system (*L* =1, red dots), for the size distribution (A) and duration distribution (B). The distributions show a clear dependence on the topology of the network. In the fully connected model (red dots), exponents are *α* = 1.48 ± 0.01 and *τ* = 2.05 ± 0.01, very close to the values characteristic of the branching model of neural dynamics, as already observed in [7]. In the 2D system (blue dots), after a small size and duration regime characterized by exponents close to one, one finds, up to the cut-off of the distributions, a scaling regime with exponents *α* = 1.29 ± 0.01 and *τ* = 1.37 ± 0.01, markedly smaller than the mean field values.

**FIG. 4.**
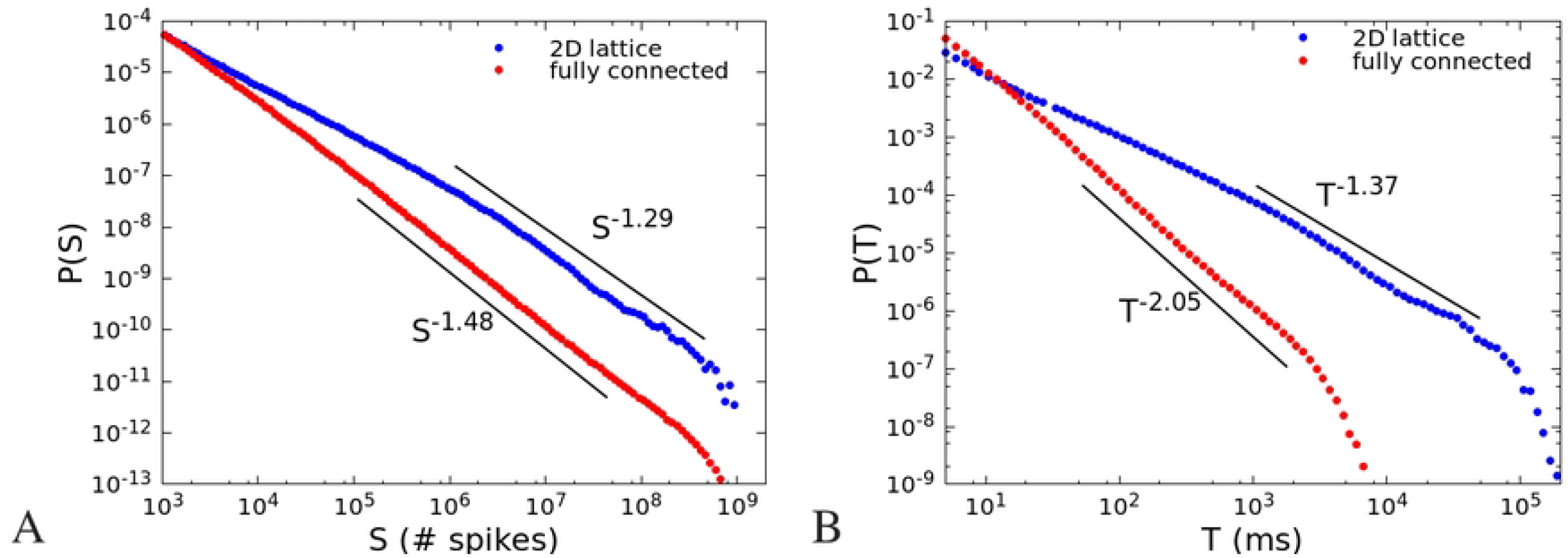
(A) Avalanche size distribution *P*(*S*) for a 2D system with *L* = 40, with a range of connections *λ* = 1 (blue dots) and for a fully connected system with *L* = 1 (red dots). Other parameters are 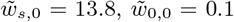, *h* = 10^-8^, *δ* = 1 ms. The number of neurons is 10^8^ per site. (B) Avalanche duration distribution *P*(*T*) for the same system and parameters. The fits of the power law exponents are done with the Python program “powerlaw” [25] in the ranges indicated by the black lines.

We have then considered the relation 〈*S*〉(*t*) between the duration of an avalanche and its mean size (Fig. 5A), for the same system and parameters of Fig. 4. We fit the measured data, in the same ranges of Fig. 4B, with a power law *T^γ^*. Also in this case, we observe a marked difference between the fully connected model (red dots), where the exponent is *γ* = 2.04 ± 0.01, and the 2D system where *γ* = 1.35 ± 0.01. Note that in both cases the expected relation [9, 10]

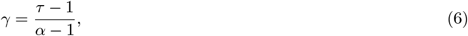

is approximately satisfied. As for the size and duration distributions, in the 2D system the scaling regime for durations 10^3^ < *T* < 5 × 10^4^ ms is preceded by a different scaling behaviour with an exponent close to 2.

**FIG. 5.**
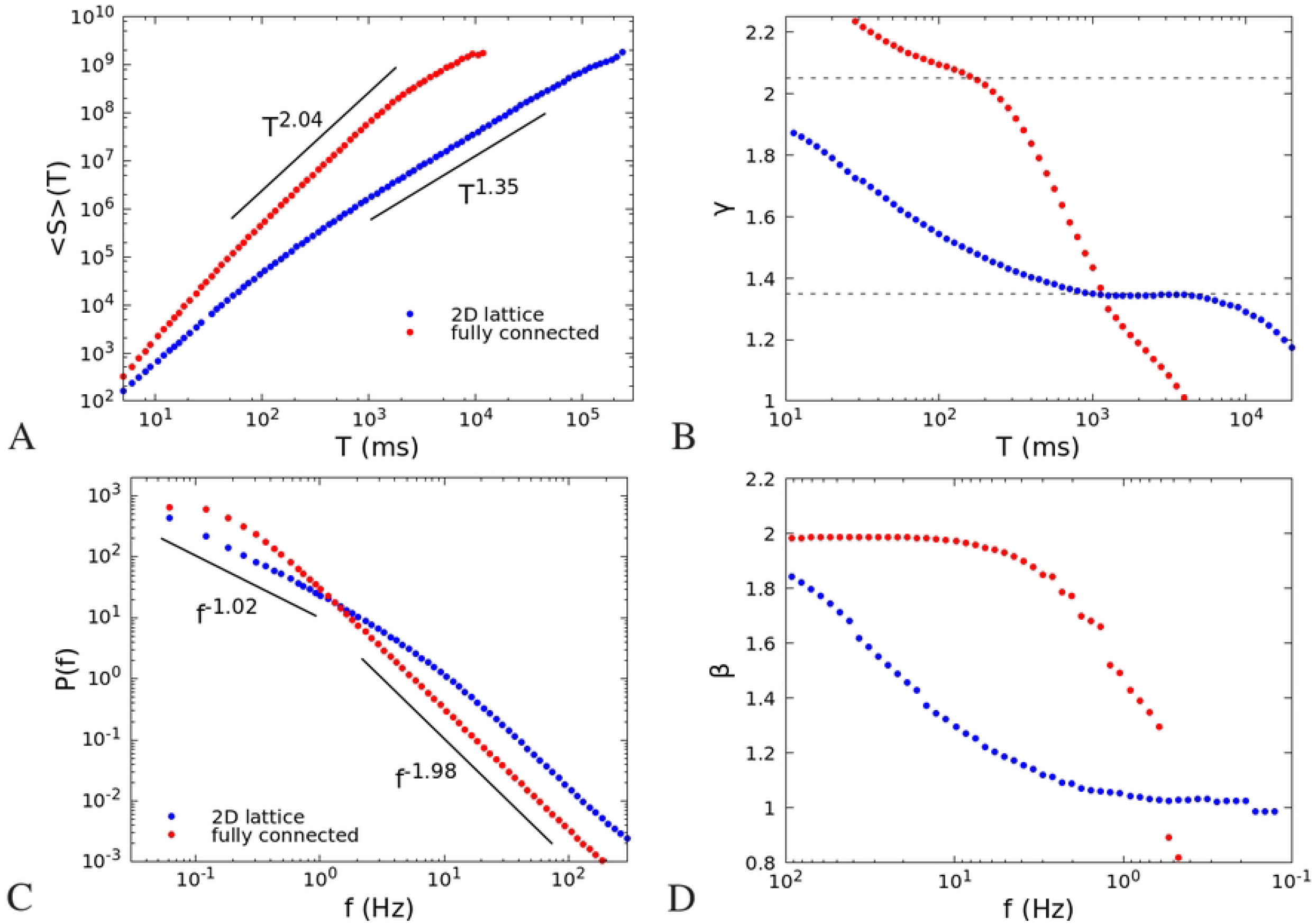
(A) Mean size of an avalanche as a function of its duration, for the same system and parameters of Fig. 4; (B) Exponent *γ* of a fit of 〈*S*〉(*T*) with a function *T^γ^*, in a sliding window [*T*, 10*T*]. The dashed lines show the exponents given by the fits of 〈*S*〉(*T*) in panel (A), and used in Fig. 6 for the collapse of the avalanche shapes. (C) Power spectrum *P*(*f*) of the single site firing rate; (D) Exponent *β* of a fit of the power spectrum *P*(*f*) with a function *f^-β^*, in a sliding window [0.1*f, f*]. To make the comparison with panel B easier, the scale on the x-axis is inverted: high frequencies (small times) are on the left, low frequencies (large times) on the right,

To put in evidence the dependence of the exponent *γ* on the range of durations chosen for the fit, in Fig. 5B we show the exponent *γ* of a fit of 〈*S*〉(*t*) with a power law *T^γ^* in a sliding window [*T*, 10*T*]. Intervals of durations where the exponent is nearly constant represent ranges where 〈*S*〉(*t*) can be well fitted by a power law. In the fully connected case (red dots) there is a range of durations, between 10^2^ and 10^3^ ms, where 〈*S*〉(*t*) ~ *T^γ^* with *γ* ≃ 2, as already observed in [7], while for longer durations the exponent drops to lower values due to the cut-off of the distribution. On the other hand, in the 2D case (blue dots), we observe a range between 10^3^ and 10^4^ where 〈*S*〉(*t*) ~ *T^γ^* with *γ* ≃ 1.3.

To investigate the relation between the exponent *γ* of 〈*S*〉(*t*) and the exponent *β* of the power spectrum, in Fig. 5C we plot the power spectrum *P*(*f*) of the single site firing rate defined in Eq. (5), for the same system size and parameters of Fig. 5A,B and Fig. 4. The power spectrum of the fully connected system is well fitted by a Lorentzian, with a white noise behaviour for frequencies lower than 1 Hz, and a decay with an exponent *β* = 1.98 ± 0.02 for frequencies larger than 1 Hz. Conversely the spectrum of the 2D system, as anticipated analitically for the *N* → ∞ case, shows a decay with an exponent *β* = 1.02 ± 0.02 for frequencies between 0.05 and 1 Hz, intermediate between white noise at lower frequencies (not shown) and brown noise at higher ones. As we have done with the exponent *γ* in Fig. 5B, in Fig. 5D we plot the exponent *β* of a fit *P*(*f*) ~ *f^-β^* in a sliding window of range [0.1*f, f*]. Comparing Fig. 5D with Fig. 5B, it can be seen that exponents *γ* and *β* have a similar dependence respectively on the avalanche duration and on the inverse frequency. In the case of the fully connected system (red dots) both exponents are around 2 in the ranges [10^2^, 10^3^] ms and [1, 10] Hz, and decay to lower values for longer avalanches or smaller frequencies. In the case of the 2D system the exponents are nearly constant in the ranges [10^3^, 10^4^] ms and [0.1, 1] Hz, although they tend to quite different values, namely *β* ≃ 1, *γ* ≃ 1.3. The reason of this discrepancy is to be further investigated.

### Scaling of the shape of avalanches

Together with the relation Eq. (6), another test of the “criticality” hypothesis for avalanche activity, is the scaling of the shape of the avalanches. Denoting with *V*(*t*) the mean number of spikes observed at time *t* during an avalanche of duration *T*, the total size of the avalanche is given by 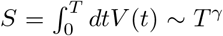, so it is expected that the normalized shape *V*(*t*)T^1–*γ*^ depends only on the rescaled time *t/T*. This relation should hold as long as the mean size 〈*S*〉(*t*) is well fitted by a power law *T^γ^*, that is for durations where the exponent of the sliding window fits are nearly constant in Fig. 5B. Looking at Fig. 5B, we expect a collapse of the shapes in the interval 10^2^ < *T* < 10^3^ ms for the fully connected system, with a value *γ* ~ 2.04, and in the interval 10^3^ < *T* < 10^4^ ms for the 2D system, with a value *γ* ~ 1.35. We highlight such values of *γ* by dashed lines in the figure. The collapse of the shapes is indeed quite well verified, for such values and duration ranges, as shown in Fig. 6A,B. Note that, as expected, an exponent *γ* ≃ 2 corresponds to a shape that is nearly parabolic (fully connected system), while in the case of the 2D system the exponent *γ* – 1.3 corresponds to a more flattened shape [26].

**FIG. 6.**
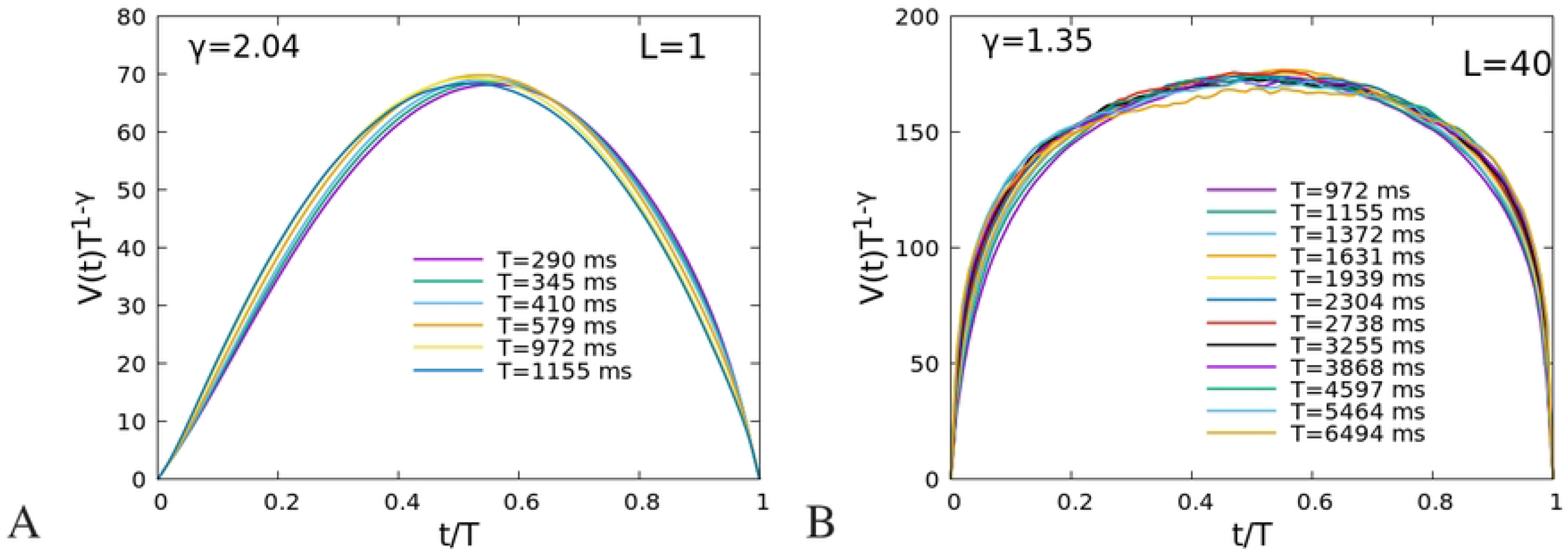
(A) Average shape of the avalanches having duration in an interval [*T*/1.09, 1.09*T*] centered on the duration *T* listed in the legend, divided by *T*^1–*γ*^, for the same parameters of Fig. 4 and for the fully connected system (*L* = 1); (B) The same for the 2D system (*L* = 40).

### Dependence on the fraction of inhibitory neurons

In this section we investigate the role of the ratio between excitatory and inhibitory neurons, by simulating the system also for a fraction 80/20 of excitatory and inhibitory neurons, more similar to cortical networks. The total number of neurons on each site of the lattice is 2 × 10^8^ as in the previously studied (50/50) case, so that we have *n_E_* = 1.6 × 10^8^ and *n_I_* = 4 × 10^7^. Note that, as it is apparent from Eq. (3), the strength of the excitatory/inhibitory synapses is inversely proportional to the number of excitatory/inhibitory neurons, so that if the number of inhibitory neurons is decreased, the strength of inhibitory connections is correspondingly increased. In this way we ensure that the critical point corresponds to the same value of 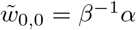 and *h* = 0.

In Fig. 7 we compare the four cases considered: fully connected 50/50, fully connected 80/20, 2D 50/50 and 2D 80/20. We note that the fraction of inhibitory neurons affects only the cut-off in the distributions *P*(*S*) and *P*(*t*), while the mean avalanche size 〈*S*〉(*t*) and the power spectrum *P*(*f*) are not affected. Moreover the cut-off has a significative change only in the 2D system, and is much less affected in the fully connected case. In the insets of Fig. 7A,B, we report a collapse of the 50/50 and 80/20 distributions in the 2D case, showing that the cutoff is approximately four (three) times smaller in the 80/20 case for the size (duration) distributions.

**FIG. 7.**
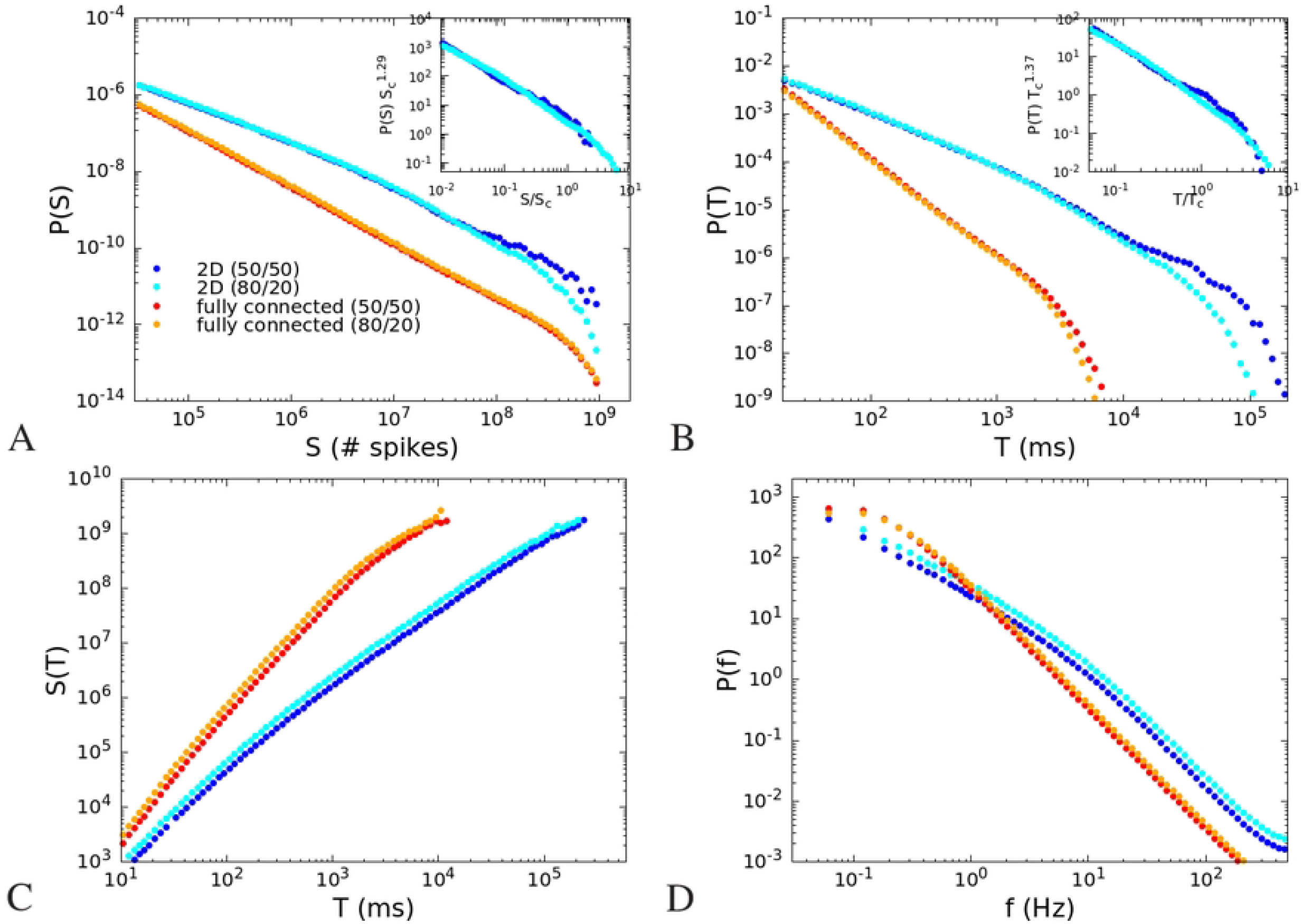
(A) Avalanche size distribution *P*(*S*) for the fully connected and 2D models, and two different fractions of excitatory and inhibitory neurons, 50/50 and 80/20. The total number of neurons on each site is 2 × 10^8^ in all cases. Other parameters as in Fig. 4 and 5. Inset: collapse of the distributions in the 2D case. The cut-off *S_c_* is 4 × 10^8^ in the 50/50 case and 10^8^ in the 80/20 case. (B) Avalanche duration distribution. Inset: collapse of the distributions in the 2D case. The cut-off *T_c_* is 3 × 10^4^ in the 50/50 case and 10^4^ in the 80/20 case. (C) Mean size of the avalanche as a function of the duration. (D) Power spectrum of the firing rate.

In conclusion, we can affirm that the observed dependence of critical exponents on the spatial dimensionality is preserved also for different fractions of inhibitory neurons.

## Conclusions

We have studied the stochastic Wilson Cowan model on a 2D lattice, with connections decaying exponentially with the distance. We use a connection weight that decays exponentially with distance to model the structural anatomical connectivity that exhibits exponential decay with distance. Recent research indeed, using data from retrograde tracer injections, shows that influence of interareal distance on connectivity patterns is conform to an exponential distance rule (EDR), according to which the projection lengths decay exponentially with inter-areal separation [27–31]. Spatial decay constant is of order of few mm for white matter, and few tenths of mm for gray matter, in mouse and macaque, [28, 29]. Notably, in the 2D model, as in the fully connected case, varying the difference between the strength of excitatory and inhibitory connections, the model undergoes a second order transition from a phase where the activity tends to zero in absence of external input, to a phase where the activity is self-sustained even in absence of external inputs. At the critical point, one of the relaxation times diverge. While the fully connected model is characterized by just two relaxation times, one of which is always lower than *α*^-1^, where *α* is the deactivation rate of neurons, the 2D system has a spectrum of relaxation times, related to the different Fourier modes that can be defined on the lattice. Such relaxation times become proportional to 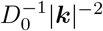 at the critical point, where *D*_0_ is a constant and, as a consequence, the model in 2D shows a logarithmic decay of the relaxation functions and a f^-1^ behaviour of the power spectrum, in an interval of frequencies between 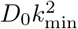 and 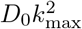, where 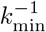, is the lattice size, and 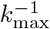 is the lattice constant. We emphasize that the 1/*f* behaviour is observed for the single site activity (or the activity of a localized group of sites), being the superposition of all the Fourier components with a spectrum of relaxation times. Conversely, the activity of the entire system corresponds to the single Fourier component ***k*** = 0 and therefore exhibits a 1/*f*^2^ behavior. Moreover, we have shown that the change in the behaviour of the power spectrum is related to a marked change in the exponents of the avalanche distributions. Note that, in the case of the 2D system, we have defined avalanches considering the activity on a single site and not in the whole system, in order to use the same signal used to define the power spectrum. This choice has been also determined by the observation that, in a large 2D system, activity in the entire system never goes to zero, which would make the introduction of a thresholding procedure necessary to define avalanches. Clearly, although avalanches are measured locally, their distribution reflects the activity of all the sites of the system. In the 2D case, the exponents *α* and *τ* of the size and duration distributions become smaller than the ones predicted in the mean field case, respectively *α* ≃ 1.3 and *τ* ≃ 1.4, while the exponent *γ* that relates the mean size to the avalanche duration decreases from 2 to 1.3. It remains to be understood the discrepancy between *γ* and the exponent *β* of the power spectrum. Notably in the 2D model the avalanche shape collapse is quite well verified with a value *γ* ≃ 1.35, showing a more flattered shape, while it is *γ* ≃ 2 in the fully connected case.

Discrepancy between the fully connected *γ* ≃ 2 prediction and the exponent *γ* ≃ 1.3 observed in some experiments [12], with a large interval of exponents *α* and *τ* of the size and duration distributions all falling on the *γ* ≃ 1.3 line, may be attributed to the effective topology of the measured activity. For example measurement able to record spiking activity of large areas of neuronal populations may reflect the structured topology of the network, while more localized measures (in highly connected areas) may be better approximated by mean-field i.e. fully-connected topology. It would be interesting to compare the corresponding scaling behavior of the power spectra of experimental activity in different topological conditions, to confirm the existence of different universality classes.

## Methods

We consider a two-dimensional lattice of *L* × *L* sites. On each site there are *n_E_* = *N_E_/L*^2^ excitatory neurons and *n_I_* = *N_I_/L*^2^ inhibitory ones. Define *m_r_* and *l_r_* respectively the number of active excitatory and inhibitory neurons on the site ***r*** = (*x, y*), with *x, y* = 0,…, *L* – 1.

In the Gaussian noise approximation, the temporal evolution of the system can be effectively described in terms of the coupled non-linear Langevin equations [32]

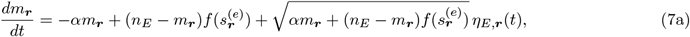

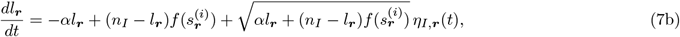

where *α* is the rate of deactivation of the neurons, *f*(*s*) is the input dependent rate of activation, 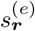 and 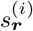 are respectively the inputs of the excitatory and inhibitory neurons on site ***r***. They are given by

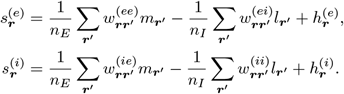

where 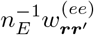 is the strength of the connection between excitatory neurons on site ***r***′ and those on site ***r***, etc… 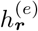 and 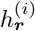 are the external inputs, *η_E,**r**_*(*t*) and *η_I,**r**_*(*t*) are Gaussian white noise functions.

The fractions of active neurons *x_**r**_* = *m_**r**_/n_E_*, *y_**r**_* = *l_**r**_/n_I_* obey therefore the equations

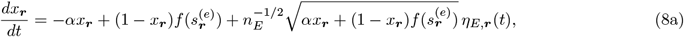

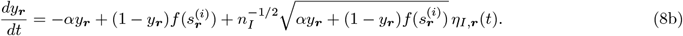

Let us suppose now that connections depend only on the distance |***r*** – ***r***′|, and the external input is independent of the site, 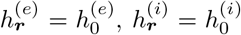. In these hypotheses the fixed point of equations (8) is the same for all sites, and corresponds to *x_**r**_* = *x*_0_, *y_**r**_* = *y*_0_, with

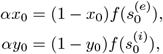

and

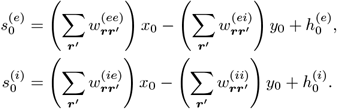

We can perform a system size expansion, changing variables from the fractions of active neurons *x_**r**_* and *y_**r**_* to the deviations of the fractions with respect to the fixed point. Defining 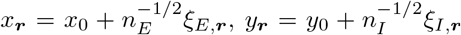, substituting in Eq. (8), and neglecting terms that are small when the number of neurons is large, we obtain the linear Langevin equations

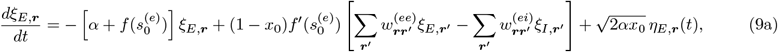

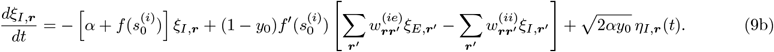

The equations (9) are translationally invariant and linear, therefore it is convenient to perform a Fourier transform and define

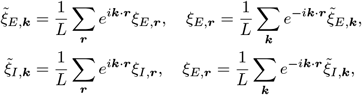

where 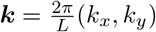, with *k_x_, k_y_* = 0,…, *L* – 1.

Substituting in (9), we obtain the decoupled equations for each of the *L* × *L* pairs of Fourier modes 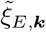 and 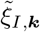,

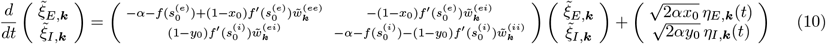

where 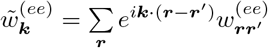, etc…, 〈*η_E,**k**_*(*t*)*η_E,**k**′_*(*t*′)〉 = *δ*_***k**′,–**k**^δ^*_(*t* – *t*′), etc…

If the connections do not depend on the post-synaptic neuron, but only on the pre-synaptic one, that is 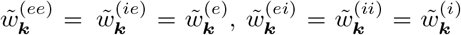, and the external input is 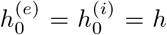, we can further simplify the equations, making the variable substitution 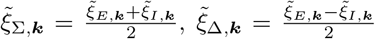. In this case, the fixed point values *x*_0_ and *y*_0_ of the excitatory and inhibitory fraction of active neurons are the same, so that we can define *x*_0_ = *y*_0_ = Σ_0_, and 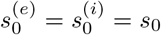, and the matrix in Eq. (10) becomes upper triangular

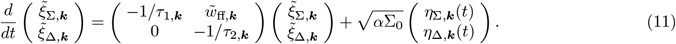

where

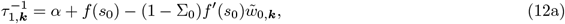

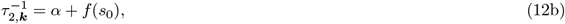

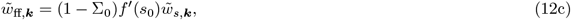

and

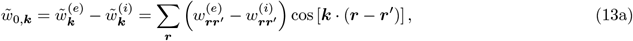

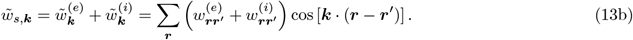

where we have used the fact that 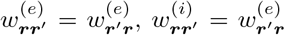. Note that *τ*_2,***k***_ is independent of ***k***. The fixed point input *s*_0_ can be written as 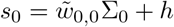, and we report here for convenience the form of the fixed point equation,

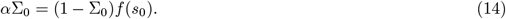

As we have seen, each of the Fourier modes behaves exactly as the total fraction of neurons in the model with all-to-all connections, but with parameters 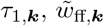 that depend on the Fourier mode ***k***. The autocorrelation function of the fluctuations 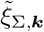 is therefore given by [22]

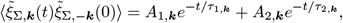

where

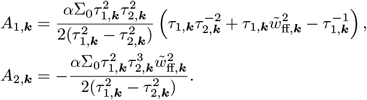

By performing an inverse Fourier transform, we can find the autocorrelation function of 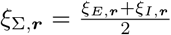, which is given by

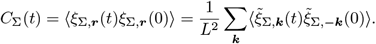

The power spectrum of the variable ξ_Σ,***r***_(*t*) is therefore given by the Wiener-Khinchin theorem as

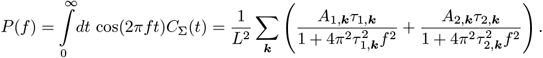

The linear equations (10) depend basically on parameters defined at the fixed point. The fixed point can undergo different kinds of transitions (bifurcations). We will consider here the case of the transcritical bifurcation, where one of the eigenvalues of the matrix in Eq. (11) vanishes, in particular we will consider the case in which the eigenvalue 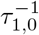 corresponding to the mode ***k*** = 0 vanishes, so that

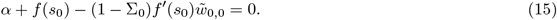

For values of |***k***| smaller than *λ*^-1^, where *λ* is the range of the connections 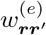, and 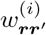, taking the first two terms of the Taylor expansion of the cosine in Eq. (13a), we have

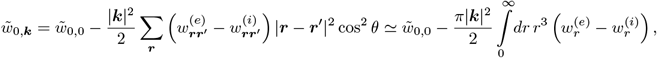

where *θ* is the angle between ***k*** and ***r*** – ***r***′. Putting this expression in Eq. (12a), and using Eq. (15), we obtain that 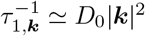, where

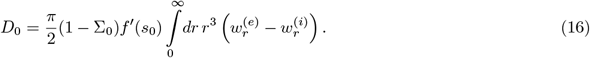

The values of *τ*_1,***k***_ therefore diverge for |***k***| → 0, so that we can consider *τ*_1,***k***_ ≫ *τ*_2,***k***_ for low values of the wave number. In this case, we can approximate

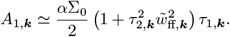

Therefore both *τ*_1,***k***_ and *A*_1,***k***_ diverge for |***k***| → 0, while *τ*_2,***k***_, *A*_2,***k***_ e 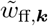 remain finite. Neglecting terms relative to *τ*_2,***k***_ in the power spectrum, we obtain

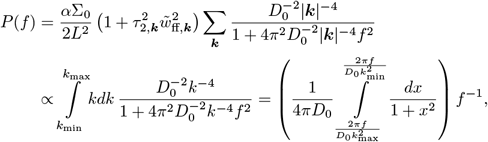

so that *P*(*f*) ∝ *f*^-1^ for 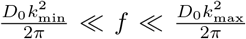, where *k_min_* and *k_max_* correspond to the reciprocal lattice size and lattice constant.

## Acknowledgments

AdC, IA and LdA acknowledge financial support from the MIUR PRIN 2017WZFTZP “Stochastic forecasting in complex systems”. AS acknowledges financial support form MIUR PRIN 201798CZLJ. LdA and AS acknowledge support from Program (VAnviteLli pEr la RicErca: VALERE) 2019 financed by the University of Campania “L. Vanvitelli”.

